# PIPENN-EMB: ensemble net and protein embeddings generalise protein interface prediction beyond homology

**DOI:** 10.1101/2024.10.31.621117

**Authors:** David P. G. Thomas, Carlos M. Garcia Fernandez, Reza Haydarlou, K. Anton Feenstra

## Abstract

Protein interactions are crucial for understanding biological functions and disease mechanisms, but predicting these remains a complex task in computational biology. Increasingly, Deep Learning models are having success in interface prediction. This study presents PIPENN-EMB which explores the added value of using embeddings from the `ProtT5-XL` protein language model. Our results show substantial improvement over the previously published PIPENN model for protein interaction interface prediction, reaching an MCC of 0.313 vs. 0.249, and AUC-ROC 0.800 vs. 0.755 on the `BIO_DL_TE` test set. We furthermore show that these embeddings cover a broad range of ‘hand-crafted’ protein features in ablation studies. PIPENN-EMB reaches state-of-the-art performance on the `ZK448` dataset for protein-protein interface prediction. We showcase predictions on 25 resistance-related proteins from *Mycobacterium tuberculosis*. Furthermore, whereas other state-of-the-art sequence-based methods perform worse for proteins that have little recognisable homology in their training data, PIPENN-EMB generalises to remote homologs, yielding stable AUC-ROC across all three test sets with less than 30% sequence identity to the training dataset, and even to proteins with less than 15% sequence identity.

**Availability:** Webserver, source code and datasets at www.ibi.vu.nl/programs/pipennemb/

## Introduction

Protein interactions are fundamental to cellular processes^1^. These interactions drive signal transduction, enzymatic activity, structural support, and the regulation of gene expression^2^. Disruptions in this mechanism can lead to various diseases^3^, making them a key focus for therapeutic interventions^4–6^. Tackling protein interactions can help understand biological functions and disease mechanisms. Despite having annotations for protein interactions in a pair-based approach, i.e. knowing if two proteins interact, most of these known interactions lack detailed structural information and thereby making them therapeutically non-viable targets^2,7^. Understanding the binding site whereby proteins interact can shed light on how protein interactions come about; residues forming the binding site are called interface residues. Ongoing research is moving forward to understanding which residues belong to the interface by turning the problem into a prediction task^8–13^.

Recent advances in machine learning approaches demonstrate strong performance in predicting different types of protein annotations^10,12^. Algorithms addressing protein interface prediction use two main approaches: structure-based and sequence-based. Structure-based methods generally outperform sequence-based ones. However, despite AlphaFold2^14^, reliable structural information is still not available for many important types of proteins and protein regions^15,16^. Moreover, the usefulness of predicted structures as input for prediction of functional properties such as interface regions or binding sites may be still quite limited^17–19^. On the other hand, sequence-based approaches, while also dependent on structure-based annotations for training, can exploit the benefits of pre-training and transfer learning from the abundantly available sequence data^20–25^. Transfer learning techniques, such as using embeddings, leverage a self-supervised learning approach to learn powerful representation of proteins at the amino acid level^20,25^. Previous work demonstrated the superior capability of embeddings when compared to ‘hand-crafted’ features for predicting protein properties^26–28^. Sequence-based models typically employ architectures developed for modeling time series or text data, such as Convolutional Neural Networks (CNN), Recurrent Neural Networks (RNN) or Transformers^29,30^. Moreover, ensemble networks are an interesting option to maximize performance^12^. These ensembles exhibit greater generalisability compared to the regular architectures, albeit at the expense of reduced interpretability^31^.

In this study, we explore two distinct enhancements to the PIPENN model^12^, our previously developed sequence-based ensemble predictor for protein interfaces. First, we retrained PIPENN using embeddings from `ProtT5-XL`^20^, using the `BioDL_P_TR` training set. Subsequently, we updated the ensemble neural network, which we will refer to as PIPENN-EMB, and evaluate its performance on the PIPENN `BioDL_P_TE` testset, and the external `ZK448` benchmark dataset. To evaluate PIPENN-EMB’s potential to predict protein interaction interface residues on a new unseen dataset, we applied it to *Mycobacterium tuberculosis* (MTB). MTB infection causes tuberculosis (TB), which continues to be a major cause of death worldwide; in 2022 approximately 1.3 million deaths according to the World Health Organisation^32^. Each year, around 300,000 people contract drug-resistant MTB. Our exploration centers on PIPENN-EMB’s potential to predict interface residues in MTB drug resistance-related proteins^33,34^. Such an analysis may contribute to our understanding of how protein interfaces may drive TB drug resistance.

## Methods

### Data Collection and Preprocessing

We used the PIPENN training data set (`BioDL_P_TR`) and its fully independent test data set (`BioDL_P_TE`), which were newly constructed by Stringer et al. (2022)^12^, and the `ZK448` benchmark data set from Zhang & Kurgan (2018)^10^. Both contain residues annotated for protein-protein interactions. In summary, `BioDL_P` was generated using annotations from protein structures from PDB, following specific annotation criteria and sequence mapping protocols. Post-processing steps, including sequence clustering at 25% and filtering on sequence length between 30–700, were applied to form non-redundant training and testing sets. For evaluating the performance of our trained models on various independent test sets, we used the *Equal* method^10^, by which a cutoff point is selected where number of actual positives is equal to the number of predicted positives.

### Model Architectures

The PIPENN architectures are composed of multiple building blocks, each containing various hyperparameters. Different compositions enable learning architectures to provide great flexibility and a broad application area. The following gives a high-level overview of the PIPENN architectures, further detail may be found in Stringer et al. (2022)^12^.

PIPENN is based on an ensemble of neural networks, which is referred to as *ensnet*. It is a neural network that uses the protein interface predictions of the following six neural architectures as input to determine whether an amino acid is part of an interface: *ann* is a fully connected architecture; *dnet* is a dilated CNN; *rnet* is a residual CNN; *rnn* consists of two layers of Gated Recurrent Unit (GRU) cells; and *cnet* is a hybrid model that combines the strengths of *rnet* and *rnn* to enhance performance. For training of PIPENN-EMB with embeddings features, we utilised the same hyperparameters as established for the PIPENN models.

### Protein Language Model Embeddings

Embeddings are a dense vector representation, capturing complex relationships and characteristics of amino acids that are not readily apparent in their raw form. Protein Language Models (PLMs) extend the concepts of Natural Language Processing (NLP) language models to the realm of protein sequences: amino acids are treated as tokens (cf. words in NLP), and entire protein sequences are regarded as sentences^20,35^.

PLMs are initially trained in a self-supervised manner, where the model learns to predict masked amino acids in known protein sequences. This training uses large datasets of protein sequences without any annotations, treating the sequences purely as text data. Here, we used the `ProtT5-XL` PLM, which was created by Elnaggar et al. (2019)^20^ using the UniRef50 database^36^. UniRef50 is approximately eight times larger than the largest datasets previously utilized for PLMs, resulting in a 5-fold increase in the number of tokens^37^. We integrated embeddings from `ProtT5-XL` in all our datasets and then measured the performance of the new PIPENN-EMB models in comparison with the previous PIPENN.

### Feature Ablation

The ablation study conducted at the feature level involved dividing the features into four categories: (1) Structural Information (SI), including secondary structure and surface accessibility; (2) Position Specific Scoring Matrix (PSSM), representing evolutionary conservation information for each sequence, generated using `PSI-BLAST`; (3) Protein Length (PL), representing the number of amino acids in a protein; (4) Embeddings (EMB), from `ProtT5-XL` as described above. In order to assess the impact of embeddings on the performance, we retrained the models using different combinations of the four feature categories. The training was performed on the PIPENN `BioDL_P_TR` data set and tested on the test data set `BioDL_P_TE`.

### Mycobacterium tuberculosis use-case

As an additional external test-set, we selected proteins with known antimicrobial resistance phenotypes in *Mycobacterium tuberculosis* (MTB) from a GWAS dataset acquired from TB-profiler^38^. Interface annotations were retrieved from the PDB-KB API^39^. We excluded proteins longer than 1000 residues, due to length constraints imposed by the prediction methods. Additionally, proteins lacking protein annotation from the PDB-KB API were also omitted from the analysis. The selected 25 proteins constitute the `MTB` test set.

### Comparison to state-of-the-art

We also used the `MTB` test set to compare the performance of our sequence-based interface prediction model (PIPENN-EMB) with other top-performing sequence-based and parameter-free structure-based prediction methods. Specifically, we evaluated against Seq-InSite^40^, CSM-potential2^41^, and PeSTo^42^.

### Homology detection

It is well known that homology is a strong source of predictions for protein properties, however, in the context of method comparison it leads to data leakage between training and test performance^43^. To investigate this beyond the usual 25% sequence identity filtering that is now regularly done^44^, we compare observed performance per protein with its similarity to the respective method’s training data. For the detection of homology between training data and protein sequences of interest, we ran `BLASTP` locally (NCBI-BLAST-2.15.0+)^45,46^. Selection of hits was based on the lowest E-value.

We obtained the training data set for CSM-potential2 from their webserver https://biosig.lab.uq.edu.au/csm_potential/data^41^. The PeSTo training data was obtained from their repository https://github.com/LBM-EPFL/PeSTo/blob/main/data/datasets/subunits_train_set.txt^42^. For Seq-InSite the training set `trainDset_without500+_Pid_Pseq_label_V10Aug2021.txt` was kindly provided by the authors^40^. To ensure that local alignments reflect a meaningful proportion of the total query sequence coverage, we employed:

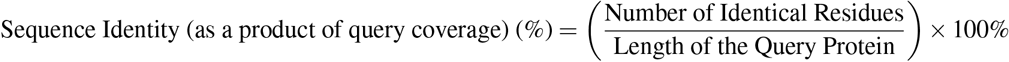

This approach enables the calculation of each hit’s coverage as a percentage of the total query length, yielding a unified metric. By correlating the predictive power of the models with overall sequence identity and coverage, we can evaluate whether the models are effectively generalising or merely identifying homologous sequences. Pearson correlation coefficient was used to evaluate relationships between sequence similarity and model performance. Correlations and p-value were calculated using the `scipy` package, considering p-values *<*0.05 statistically significant.

## Results

To corroborate the strength of embeddings as input features compared to hand-crafted features, for protein interface prediction, we constructed several neural network models with and without embedding features. All models were trained on the protein-protein training set `BioDL_P_TR`. Evaluation and selection of models were done on a 20% subset of the training set. Final selected models were subsequently tested on the `ZK448`, `BioDL_P_TE`, and `MTB` datasets, as detailed below.

### Embeddings yield higher performance

We studied the improvement achieved after introducing `ProtT5-XL` embeddings as input features for the ensemble network (*ensnet*) in the `BioDL_P_TE`. Substantial improvement can be seen for all metrics when using the embeddings, as shown inTable 1. To further discern the primary contributors to model performance, we examined various combinations of features: Protein Length (PL), Structural Information (SI), Profiles (PSSM) and `ProtT5-XL` Embeddings (EMB). We evaluated EMB, EMB+PL, EMB+SI, EMB+SI+PSSM and EMB+SI+ PSSM+PL (Table 1). For all combinations, differences were minimal, with MCC ranging from 0.290 for EMB+PSSM to 0.313 for EMB+PL. None of the ‘hand-crafted’ features, SI and PSSM, demonstrate any improvement. These results align with previous findings on the strengths of embeddings^26^. From a practical standpoint, this implies that the computation of hand-crafted features has become unnecessary.

**Table 1.** Impact of embeddings and other feature groups on the performance of the ensemble predictors.

| Features | F1 | MCC | AP | AUC | ACC | SPEC |
| --- | --- | --- | --- | --- | --- | --- |
| 1H+SI+PSSM+PL | 0.339 | 0.249 | 0.302 | 0.755 | 0.840 | 0.909 |
| EMB | **0.396** | **0.313** | 0.372 | **0.798** | **0.854** | **0.917** |
| EMB+PL | **0.395** | **0.313** | **0.377** | **0.800** | **0.854** | **0.917** |
| EMB+SI | 0.373 | 0.287 | 0.329 | 0.772 | 0.849 | **0.914** |
| EMB+SI+PSSM | 0.384 | 0.299 | 0.361 | 0.793 | 0.851 | **0.915** |
| EMB+PL+SI+PSSM | 0.382 | 0.290 | 0.321 | 0.784 | **0.858** | **0.913** |

Feature ablation study of *ensnet* model without (top) and with (EMB) embedding features, trained on `BioDL_P_TR` and tested on `BioDL_P_TE`. Reported metrics are Accuracy (ACC), Specificity (SPEC), F1 Score, Matthews Correlation Coefficient (MCC), Average Precision (AP), and Area Under the ROC Curve (AUC). Scores up to 0.005 below highest per metric in bold.

### Composition of the ensemble network matters

Our ensemble network is a flexible neural network. Given that it orchestrates the other six PIPENN architectures and utilises their predictions to perform its own predictions, we decided to assess the contribution of each architecture to the ensemble model. Table 2 shows the performance of various combinations of the individual PIPENN architectures. Each combination comprises an ensemble network. Interestingly, we observed that a combination of *rnn, rnet*, and *unet* exhibits slightly superior performance compared to other combinations. While this improvement may seem marginal, it holds significance from a practical standpoint, as it allows for simplification of the overall ensemble architecture. Moreover, excluding *ann, cnet*, and *dnet* from the ensemble results in a reduction of approximately 6 million parameters.

**Table 2.** Performance results of ensemble models comprising different architecture combinations.

| ann | cnet | dnet | rnn | rnet | unet | F1 | MCC | AP | AUC | ACC | SPEC |
| --- | --- | --- | --- | --- | --- | --- | --- | --- | --- | --- | --- |
| – | – | – | ✓ | ✓ | ✓ | **0.395** | **0.313** | **0.377** | **0.800** | **0.854** | **0.917** |
| – | ✓ | ✓ | ✓ | ✓ | ✓ | 0.388 | 0.294 | 0.362 | 0.788 | **0.851** | 0.890 |
| – | ✓ | – | ✓ | ✓ | – | 0.372 | 0.302 | 0.360 | 0.788 | **0.855** | 0.886 |
| – | – | – | – | ✓ | ✓ | 0.385 | 0.294 | 0.364 | 0.796 | **0.854** | 0.901 |
| ✓ | ✓ | ✓ | ✓ | ✓ | ✓ | 0.365 | 0.287 | 0.352 | 0.787 | 0.843 | 0.895 |

Selected combinations per row, trained on BiODL\_P\_TR and tested on BiODL\_P\_TE. Metrics as in Table 1, up to 0.005 below highest in bold.

We designate the suite of *rnn, rnet*, and *unet* neural net models along with the *ensnet* ensemble net, incorporating the EMB+PL features, as PIPENN-EMB. We will further explore the performance of the PIPENN-EMB *ensnet* model.

### Comparison to state-of-the-art methods

To put the improved PIPENN-EMB performance in context, we compared it against leading sequence-based protein-protein interface prediction methods on the benchmark `ZK448` dataset. Table 3 confirms that PIPENN-EMB significantly surpasses the previous PIPENN model across all metrics except accuracy. Other recent methods such as DELPHI^47^ and EnsemPPIs^29^ are somewhat inbetween. The very recent Seq-InSite seems to stand out, and according to its authors even exceeds structure-based predictions^40^. Only for the Seq-InSite method we were able to obtain prediction scores on the `BioDL_P_TE`, `ZK448` and `MTB` datasets that allows us to draw ROC and P/R plots for detailed comparison. Figure 1 shows that, consistently for both ROC and P/R, Seq-InSite outperforms PIPENN-EMB.

**Table 3.** Comparing PIPENN-EMB to state-of-the-art sequence-based predictors on the ZK448 benchmark dataset.

| Method | F1 | MCC | AP | AUC | ACC |
| --- | --- | --- | --- | --- | --- |
| Seq-InSite<sup>40</sup> | **0.535** | **0.462** | **0.617** | **0.853** | **0.874** |
| PIPENN-EMB* | 0.505 | 0.392 | 0.513 | 0.805 | 0.815 |
| EnsemPPIS<sup>29</sup> | 0.385 | 0.291 | 0.354 | 0.770 | 0.821 |
| DELPHI<sup>47</sup> | 0.364 | 0.278 | 0.326 | 0.746 | 0.848 |
| PIPENN<sup>12</sup> | 0.339 | 0.249 | 0.302 | 0.755 | 0.840 |
| SCRIBER<sup>48</sup> | 0.333 | 0.230 | 0.287 | 0.715 | n.a. |
| SPRINGS<sup>49</sup> | 0.229 | 0.111 | 0.201 | 0.625 | n.a. |

Metrics cf. Table 1. \* this work.

**Figure 1.**
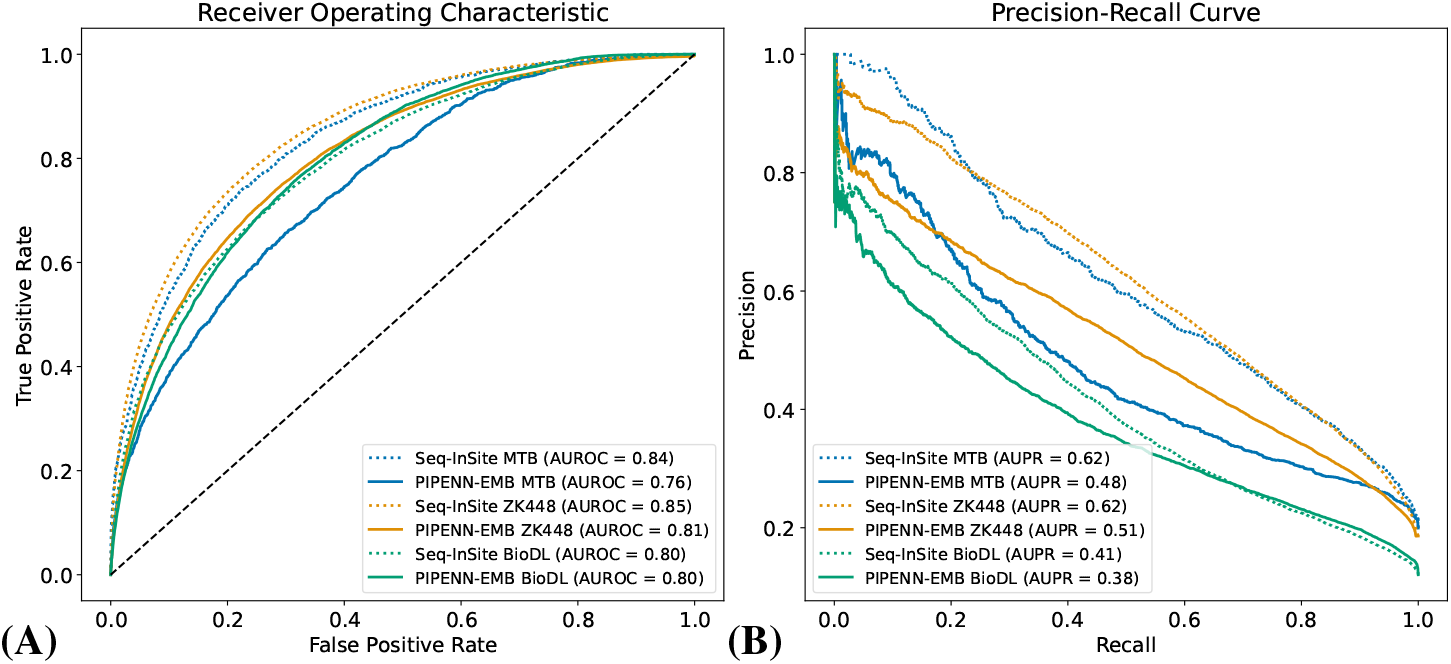
Model performance of state of the art sequence based prediction methods. (A) AUROC and (B) Precision-Recall plots for comparison of models tested on the `BioDL_P_TE` test set, the `MTB` use case, and the `ZK448` benchmark dataset. Curves correspond to the interface prediction performance for the sequence-based PIPENN-EMB and Seq-InSite. Legends report AUROC and AUPR values as measure of overall prediction capabilities.

### Sequence- and structure-based models perform on par on Mycobacterium tuberculosis proteins

Protein-protein interactions in MTB have been suggested as a key factor in driving anti-microbial resistance in TB^50^. To enhance our understanding of these interactions, we evaluate the performance of PIPENN-EMB and selected state-of-the-art methods on this specialised external use-case. This allows for further assessment of the methods’ robustness in recognising a particular subset of prokaryotic proteins, thereby providing insights into the models’ performances within more specialised scenarios and real life applications.

For comparison, we included sequence-based Seq-InSite^40^, and structure-based CSM-Potential2 [41, CSM] and PeSTo^42^. Table 4 shows that sequence-based PIPENN-EMB performing overall quite close with the structure-based predictors, with an AUC score of 0.659, compared to 0.754 for structure-based PeSTo, and 0.623 for CSM, when using AlphaFold2 (AF2) structures. With an MCC of 0.319, PIPENN-EMB performs inbetween both structure-based methods, resp. 0.463 and 0.244. The very recent sequence-based Seq-InSite can be seen to outperform the other methods, including PIPENN-EMB, on both AUC and AP (resp. 0.731 and 0.415). The AP of PIPENN-EMB was 0.320, underscoring its capability to accurately extract critical and generalisable information from sequence characteristics to predict protein interfaces.

**Table 4.**
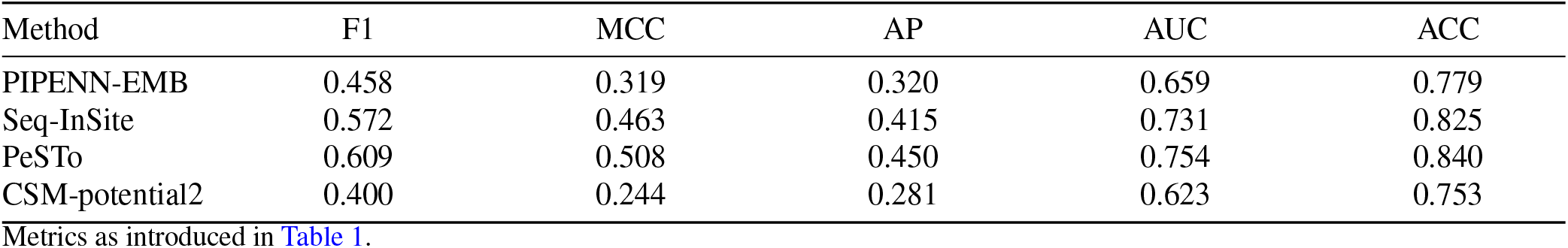
Comparing state-of-the-art sequence and structure based prediction methods on the MTB use-case dataset.

| Method | F1 | MCC | AP | AUC | ACC |
| --- | --- | --- | --- | --- | --- |
| PIPENN-EMB | 0.458 | 0.319 | 0.320 | 0.659 | 0.779 |
| Seq-InSite | 0.572 | 0.463 | 0.415 | 0.731 | 0.825 |
| PeSTo | 0.609 | 0.508 | 0.450 | 0.754 | 0.840 |
| CSM-potential2 | 0.400 | 0.244 | 0.281 | 0.623 | 0.753 |

Metrics as introduced in Table 1.

The `MTB` dataset moreover allows us to assess generalisability of the methods tested. Since its introduction in 2018^10^, the `ZK448` dataset has been extensively used for method development, potentially introducing a bias towards this specific dataset. This raises concerns that supposedly unseen benchmark data, i.e. `ZK448`, might have influenced design decisions for methods benchmarked on it. Below, we will further explore these differences in generalising capabilities.

### Generalising beyond homology

Observed differences in performance may be influenced by homology between training and test datasets, leading to an inflated estimate of the method’s capability to generalise to new, unseen data.

To assess the dependency of models’ predictions on homology between training and test datasets, we (1) utilise `BLASTp` to find the closest hits, (2) measure the degree of sequence similarity, and (3) calculate the correlation between the degree of sequence similarity and a model’s AUC-ROC. For robustness, we performed this analysis on all three test-sets: `BioDL_P_TE`, `ZK448` and `MTB`.

The results shown significant correlations between sequence similarity and coverage to the training set and the performance of Seq-InSite in Figure 2A & B, while PIPENN-EMB shows no significant correlation on any of the test-sets in Figure 2. The intersection of the correlation trends between PIPENN-EMB and Seq-InSite suggests PIPENN-EMB’s capacity for generalisation beyond homology, contrasting with Seq-InSite’s apparent dependency on homologous sequences for performance in Figure 2A & B. Thus, the higher performance of Seq-InSite seen in Table 4 might be inflated through homology. The results emphasise the robustness of PIPENN-EMB against data leakage and its capability to maintain consistent performance irrespective of sequence similarity.

**Figure 2.**
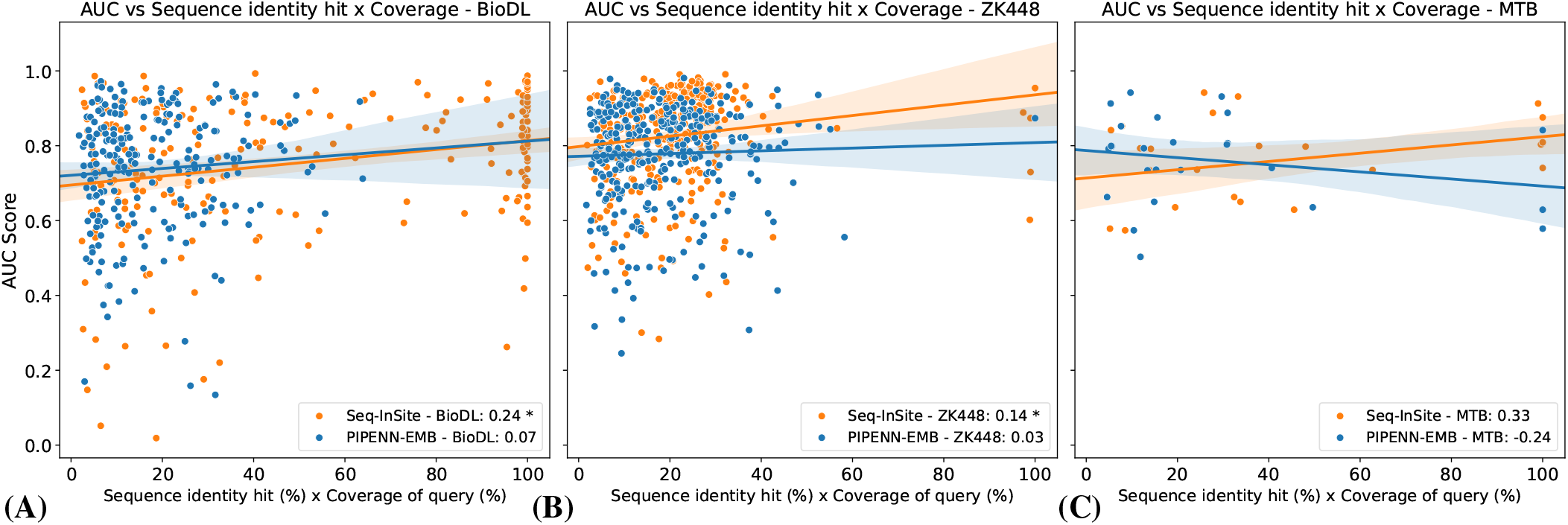
Correlation between model performance and sequence homology for different test datasets. Data points represent individual proteins from the different datasets, showing the method prediction performance in AUC-ROC versus the (the product of) sequence similarity and query coverage for hits in the method’s training set. The linear regression lines indicate trends of AUC-ROC with increasing sequence similarity and coverage. Shaded areas show the 95% confidence intervals. Correlation coefficients (R^2^) are annotated in the legends for each model, with an asterisk (*) for a p-value *<* 0.05. Results are shown for three datasets: (A) `BioDL_P_TE`, (B) `ZK448`, and (C) `MTB`.

To provide some more insight into each model’s ability to generalise predictions of proteins beyond homology with respect to the training set, we categorise proteins from these datasets into close (31-45%) and distant homologs (16-30%) and sequences that are less than 15% similar. Figure 3 shows the distribution of AUC-ROC performance of the two sequence-based methods for these three categories. For Seq-InSite, there is a noticeable drop in performance towards the more remote catagories, which is most pronounced in the `BioDL_P_TE` and `ZK448` data sets: highest scores are found in the close (31-45%) and distant category (16-30%), with a decline in performance for similarities below 15%. This pattern is indicative of Seq-InSite’s reliance on homology for predictive power, which may also be observed in Figure 2. Conversely, PIPENN-EMB displays robust performance across different levels of homology, maintaining consistent average AUC scores that do not significantly differ between categories. This consistency is evident across all three datasets, showcasing PIPENN-EMB’s ability to generalize well beyond homology. This suggests that PIPENN-EMB is particularly effective in handling diverse protein sequences, reinforcing its capabilities for predicting protein interfaces of proteins that are distant homologs from the training data.

**Figure 3.**
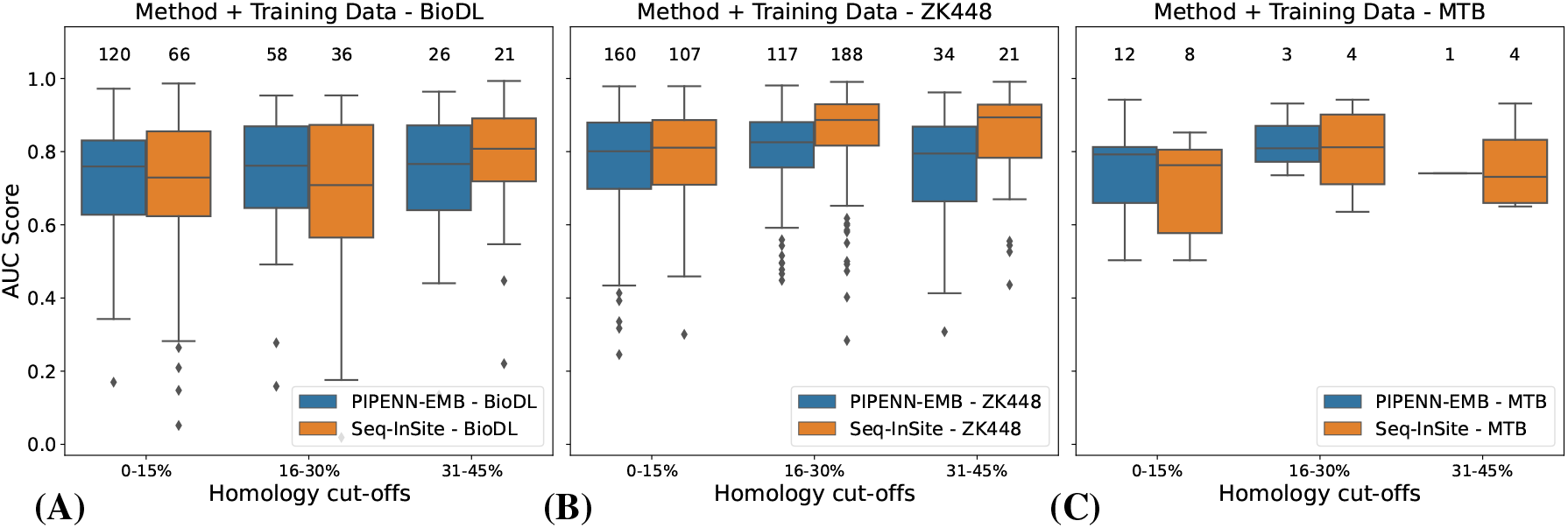
Performance of PIPENN-EMB and Seq-Insite for distant homologs from their respective training data. Boxplots illustrate the differences in the distribution of AUC scores for different homology cut-offs to the respective training data of the prediction methods. Three thresholds based on sequence identity times query coverage were applied: proteins with 0-15% identity times coverage are considered remote homologs, 16-30% are considered distant homologs, and proteins between 31-45% are considered close homologs to the respective training data for the model. The number of proteins used for each distribution is indicated above the boxes. Results are shown for three datasets: (A) `BioDL_P_TE`, (B) `ZK448`, and `MTB`.

### The Webserver

We also make PIPENN_EMB available as a webserver at www.ibi.vu.nl/programs/pipennemb/. The webserver is simple to use and is aimed at non-experts in both academia and industry. The input is only the protein sequence. The predictions are based on the DNET architecture predictions. We are still working on a version that includes also the ensemble model predictions. Expected runtimes are one to two minutes per protein, including feature generation (actual throughput may be lower due to queue time). For comparison, approximate runtimes for the PIPENN webserver are 5-10 minutes, Seq-InSite 20 minutes, CSM 5-10 minutes, and PeSto 10 seconds per protein.

## Discussion

Protein interaction interface prediction remains a challenging task despite the improvements in deep learning models that we have been seeing during the last years^12,29,40,44^. In this work we built on top of PIPENN by introducing embeddings next to a new ensemble combination achieving a comptetent performance in the benchmark `ZK448` dataset, and showcase the potential for generalisation on the new `MTB` dataset.

Embeddings provide a powerful framework to improve protein-based models by utilizing them as features^26,51,52^. Although for protein interface prediction performance seems to be reaching a plateau, these representation models are still being upgraded and show increased capability to encode complex relationships across protein sequence space. Additionally, previous work has shown that deep learning models struggle to reach generalisation in protein interface prediction due to biases in the training set and data leakage to the test set^43^. Strategies to overcome these problems lie in using more robust architectures that aim to achieve a better generalisation^31^, such as ensemble architectures, next to using smarter approaches when curating the training set^43^. A practical approach is to include new unseen data when testing new methods.

Training on structures is becoming more popular since AlphaFold2 structures are available^53^. Having access to the 3D information allows models to understand geometrical conformations involved in interactions. However, a certain degree of bias can be introduced due to the annotations that are made based on PDB structures. On top of this, representation models benefit more from sequences because there are more protein sequences available than structures. Models that combine both structural and sequential aspects of proteins are showing promising results, however, they are still in early development^54^.

The tuberculosis use-case gives insights into how PIPENN-EMB performs in identifying protein interactions within MTB resistance proteins, which includes new unseen proteins that are not homologous to the training data. Here, PIPENN-EMB outperforms state-of-the-art sequence-based Seq-InSite, particularly for very distant proteins. We verified this trend in the `BioDL_P_TE` and `ZK448` datasets, where we find it even more pronounced. This demonstrates the ability of the PIPENN-EMB model to generalise beyond homology, making it a valuable tool for predicting interactions in proteins less represented in training datasets. Moreover, novel pathogen genomes often contain unknown proteins for which no known homologs are available. Thus, the robustness to generalise interface predictions beyond homology is crucial for advancing our understanding of resistance mechanisms and to help developing new treatments.

In summary, we have contributed with the following:

(1) introducing embeddings to boost PIPENN performance;
introducing a new ensemble architecture (PIPENN-EMB) that achieves state-of-the-art performance;
(2) comparing the performance of PIPENN-EMB with other state-of-the-art sequence-based and structured-based models on the `ZK448` and `MTB` data sets, respectively; and
(3) evaluating performance biases of PIPENN-EMB and another state-of-the-art sequence-based model on the `BioDL_P_TE`, `ZK448`, and `MTB` datasets, introduced by data leakage through homology.

## Acknowledgements

We thank Jody Phelan and Nick Furnham for their insightful discussions about the role of protein interactions in the development of MTB drug resistance, and their suggestion to investigate the prediction of the binding residues involved. We also thank SeyedMohsen Hosseini and Lucian Ilie for very kindly providing the training data for their Seq-InSite method, as well as help with technical issues on their webserver.

## Author contributions statement

R.H. and K.A.F conceptualised the research goals and aims. R.H. and C.M.G.F. developed the methodology and software.

C.M.G.F. and D.P.G.T. conducted the experiments, analysed the raw results, and visualised the data presentation. All authors analysed results, and wrote and edited the manuscript.

## Additional information

The `MTB` dataset is available at https://github.com/jodyphelan/TBProfiler/blob/master/db/tbdb.bed. The training and test sets and code for training and testing are available from the PIPENN github https://github.com/ibivu/pipenn/.

